# Resistance training rejuvenates the mitochondrial methylome in aged human skeletal muscle

**DOI:** 10.1101/2021.04.13.439651

**Authors:** Bradley A. Ruple, Joshua S. Godwin, Paulo H. C. Mesquita, Shelby C. Osburn, Christopher G. Vann, Donald A. Lamb, Casey L. Sexton, Darren G. Candow, Scott C. Forbes, Andrew D. Frugé, Andreas N. Kavazis, Kaelin C. Young, Robert A. Seaborne, Adam P. Sharples, Michael D. Roberts

## Abstract

Resistance training (RT) alters skeletal muscle nuclear DNA methylation patterns (or the methylome). However, no study has examined if RT affects the mitochondrial DNA (mtDNA) methylome. Herein, ten older untrained males (65±7 years old) performed six weeks of full-body RT (twice weekly). Body composition and knee extensor torque were assessed prior to and 72 hours following the last RT session. Vastus lateralis (VL) biopsies were also obtained. VL DNA was subjected to reduced representation bisulfite sequencing providing excellent coverage across the ~16-kilobase mtDNA methylome (254 CpG sites). Various biochemical assays were also performed, and older male data were compared to younger trained males (22±2 years old, n=7). RT increased whole-body lean tissue mass (p=0.017), VL thickness (p=0.012), and knee extensor torque (p=0.029) in older males. RT also profoundly affected the mtDNA methylome in older males, as 63% (159/254) of the CpG sites demonstrated reduced methylation (p<0.05). Notably, several mtDNA sites presented a more “youthful” signature after RT in older males when comparisons were made to younger males. The 1.12 kilobase D-loop/control region on mtDNA, which regulates mtDNA replication and transcription, possessed enriched hypomethylation in older males following RT. Enhanced expression of mitochondrial H- and L-strand genes and increases in mitochondrial complex III and IV protein levels were also observed (p<0.05). This is the first study to show RT alters the mtDNA methylome in skeletal muscle. Observed methylome alterations may enhance mitochondrial transcription, and RT remarkably evokes mitochondrial methylome profiles to mimic a more youthful signature in older males.

## INTRODUCTION

Resistance training increases strength and muscle mass, and these adaptations have been attributed to various mechanisms (e.g., an increase in satellite cell number, ribosome density, etc.). Critically, molecular adaptations acutely induced by exercise precede well-documented adaptations that occur with chronic training. In this regard, several reports have noted that single exercise bouts transiently orchestrate the up- and down-regulation of hundreds of mRNA transcripts in skeletal muscle (reviewed in (Pillon et al., 2020)). These transcriptional events are complex and involve the coordinated actions of histone-modifying enzymes, transcription factors, transcriptional co-activators, and one of three RNA polymerase enzymes.

DNA methylation is a critical mechanism that regulates mRNA transcription (Eden & Cedar, 1994). This process involves a methyl group being transferred to the C-5 position of the cytosine ring, with >98% of methylation occurring at cytosine guanine dinucleotide pairing sites (i.e., CpG sites). DNA methylation is facilitated by DNA methyltransferase (DNMT) enzymes (Jurkowska, Jurkowski, & Jeltsch, 2011), and increased methylation levels in a promoter or enhancer region negatively affect mRNA transcription by either: i) impairing transcription factor binding, and/or ii) compacting DNA and making it transcriptionally inaccessible. Recent enthusiasm has surrounded how exercise alters the collective DNA methylome in skeletal muscle (Seaborne & Sharples, 2020). Barres et al. (Barres et al., 2012) provided the first evidence, at the candidate gene level, to suggest alterations in DNA methylation across canonical metabolic genes in skeletal muscle can occur within hours of a single high-intensity aerobic exercise session. Moreover, the changes in methylation inversely correlated with mRNA expression in the corresponding genes. Subsequently, novel genome-wide methylation (methylome) studies in human skeletal muscle have demonstrated that resistance exercise training (Seaborne, Strauss, Cocks, Shepherd, O’Brien, Someren, et al., 2018; Seaborne, Strauss, Cocks, Shepherd, O’Brien, van Someren, et al., 2018) and acute high intensity running exercise (Maasar et al., 2021) elicit DNA hypomethylation and the upregulation of genes related to actin/cytoskeletal, extracellular matrix, growth-related pathways, and/or metabolic pathways. These same studies have also shown that, following an earlier period of resistance training and detraining, skeletal muscle demonstrates a heightened level of hypomethylation. Importantly, some genes retain a hypomethylated signature following training-induced hypertrophy, even during a period of detraining as muscle mass returned to pre-training levels. Moreover, these genes were ‘enhanced’ during retraining as a consequence of earlier training, suggesting human skeletal muscle possesses an epigenetic memory of earlier exercise (or ‘epi-memory’) (Sharples, Stewart, & Seaborne, 2016). The biological process of aging seems to have the opposite effect on the skeletal muscle DNA methylome whereby hypermethylation seemingly accumulates (Turner et al., 2020; S. Voisin et al., 2020; S Voisin et al., 2020; Zykovich et al., 2014). However, increased physical activity (Turner et al., 2020) and resistance exercise (Blocquiaux et al., 2020) have been shown to reverse hypermethylated profiles with age to more hypomethylated signatures.

Although resistance training clearly affects molecular mechanisms related to skeletal muscle hypertrophy, the effects of resistance training on mitochondrial adaptations are less clear. Studies in younger adult populations have reported that markers indicative of mitochondrial volume increase, decrease, or do not change in response to several weeks of resistance training (Groennebaek & Vissing, 2017; Parry, Roberts, & Kavazis, 2020). We recently reported that 10 weeks of resistance training doubles skeletal muscle citrate synthase activity (a surrogate of mitochondrial volume) in older adults (Lamb, Moore, Mesquita, et al., 2020). Others have also reported that markers reflective of improved mitochondrial function occur in older adults after 14 weeks of resistance training (Parise, Brose, & Tarnopolsky, 2005). Thus, it is plausible that increases in mitochondrial biogenesis and improvements in mitochondrial function may occur in an age-dependent fashion where robust effects are more evident in older versus younger adults. Just as with the nuclear genome, the mitochondrial genome can undergo dynamic DNA methylation and demethylation (Manev, Dzitoyeva, & Chen, 2012). Earlier research in this area suggested alterations in mitochondrial DNA (mtDNA) methylation was relatively low in comparison to the dynamic changes that occur with nuclear DNA methylation (Nass, 1973). Nonetheless, other studies have since suggested that the mtDNA methylome can be transiently modulated through various perturbations. For instance, Wong et al. (Wong, Gertz, Chestnut, & Martin, 2013) used DNA pyrosequencing to demonstrate that mtDNA methylation patterns and mitochondrial DNMT3a levels are abnormal in the skeletal muscle and spinal cord of transgenic mice that develop amyotrophic lateral sclerosis. Moreover, Patil et al. (Patil et al., 2019) recently demonstrated that mtDNA methylation patterns differed in cancerous versus non-cancerous human cell lines. However, given the infancy of this research, it is unclear as to how changes in mtDNA methylation affect mitochondrial physiology.

In spite of the tremendous discoveries mentioned above in relation to the nuclear methylome and exercise training, no studies to date have examined how exercise training affects mtDNA methylation patterns in skeletal muscle. This lack of data is, in part, due to methylome studies undertaking array profiling of CpG methylation. Alternatively stated, few human exercise studies have undertaken bisulfite sequencing of skeletal muscle that allows in-depth analysis of mtDNA methylation patterns. Therefore, the current study contained multiple objectives. First, we used a genome-wide DNA bisulfite sequencing strategy (Reduced Representation Bisulfite Sequencing, or RRBS) to determine how six weeks of resistance training altered mtDNA methylation patterns in skeletal muscle of older, previously untrained males. Notably, younger resistance-trained males were also included in this analysis as a comparator group. Next, we determined if the alterations observed at the mtDNA methylome level were associated with corresponding changes in mitochondrial gene expression as well as mitochondrial protein complexes and citrate synthase activity (a marker of mitochondrial volume). Finally, we sought to determine if resistance training was able to rejuvenate the mtDNA methylome profiles of older males to profiles observed in younger males.

## RESULTS

### Training adaptations

General training adaptations in older males and a comparison to younger males are presented in Table 1. At the end of the six weeks of training, lean/soft tissue mass, vastus lateralis thickness, knee extensor peak torque and mCSA increased in older males (P = 0.017, P = 0.012, P = 0.029, and P = 0.057 respectively). However, post-training values in older males were still significantly lower than values in younger, trained males (P < 0.05 for all variables).

**Table 1.** General resistance training adaptations in older males and comparison to younger males

| Variable (units) | Mean ± SD | |
| --- | --- | --- |
| DXA lean/soft tissue mass (kg) | Older |  |
| | Pre | 58.1 ± 5.9<sup>#</sup> |
| | Post | 58.8 ± 6.2<sup>*,#</sup> |
| | Younger, trained | 64.9 ± 4.7 |
| VL muscle thickness (cm) | Older |  |
| | Pre | 2.09 ± 0.37<sup>#</sup> |
| | Post | 2.25 ± 0.27<sup>*,#</sup> |
| | Younger, trained | 3.02±0.35 |
| pQCT mCSA (cm<sup>2</sup>) | Older |  |
| | Pre | 144.5 ± 20.6<sup>#</sup> |
| | Post | 148.9 ± 19.5<sup>#</sup> |
| | Younger, trained | 194.0 ± 22.8 |
| VL peak knee extensor torque (N•m) | Older |  |
| | Pre | 144.9 ± 58.0<sup>#</sup> |
| | Post | 168.7 ± 47.3<sup>*,#</sup> |
| | Younger, trained | 223.6 ± 27.8 |

Legend: Data are from n=10 older males (age = 65 ± 7 years old) prior to and following six weeks of training as well as basal values in n=7 younger males (22 ± 2 years old) that were resistance-trained (self-reported training age of 5 ± 1 years). Abbreviations: DXA, dual-energy x-ray absorptiometry; VL, vastus lateralis. Symbols: \*, indicates increase from Pre to Post in older participants (p<0.05); #, indicates different from younger, trained males (p<0.05).

### Mitochondrial DNA methylation

The RRBS data set spanned 254 individual CpG sites mapping to the ~16 kb mtDNA region of the human genome. Comparative analysis in older participants prior to and following resistance training shows that 63% of these CpG sites (159/254) demonstrated a significant reduction in methylation following training (FDR < 0.05; change ≥ 3%; Suppl. File 1A). Even at a more stringent pre-to-post training change (≥ 5%), a large number of sites (~40%, 98/254 CpGs) possessed a hypomethylated signature. Interestingly, with the same significance criteria, no CpG sites increased in methylation following training. These collective data indicated a globally hypomethylated mtDNA profile following resistance training in older participants (P = 0.0039; Figure 1A). Human mtDNA methylation levels have previously been suggested to be dependent on sequencing coverage biases (Mechta, Ingerslev, Fabre, Picard, & Barres, 2017). We clearly show no differential read coverage issues between time points (Figure 1B), and read density had no association with methylation levels in these conditions (Figure 1C).

**Figure 1.**
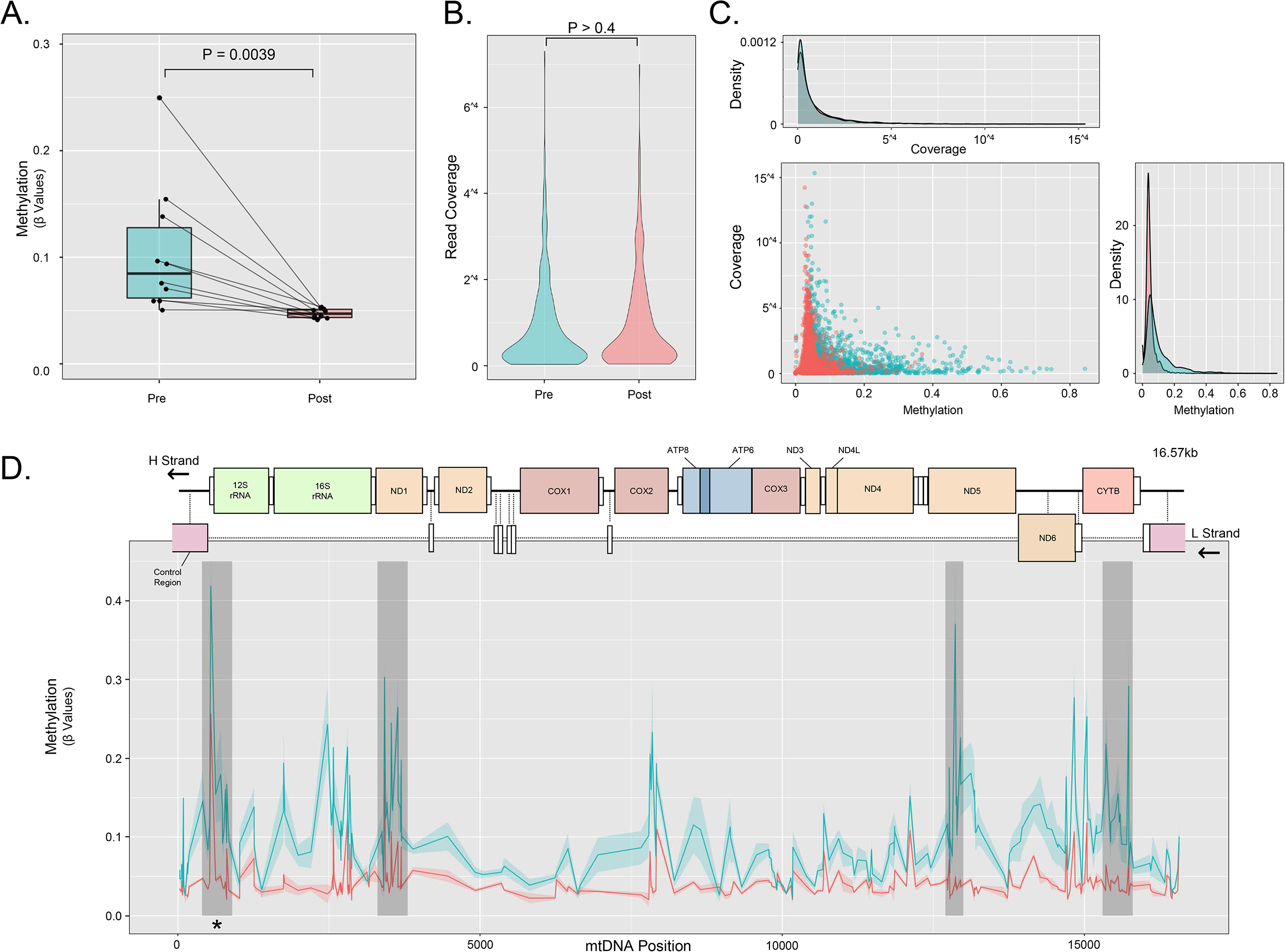
mtDNA methylome data from older males prior to and following resistance training. Legend: Data represents differential methylation in older males following six weeks of resistance training. Global mtDNA methylation significantly decreased following resistance training in older males (panel A), which we show is unlikely due to any observed differences in read coverage and association with CpG read density (panels B and C). Plotting methylation across the ~16 kb mtDNA genome (panel D), we show clear differences in methylation between two conditions (pre training, blue line; post training, red line). The four significant differentially methylated regions are highlighted in grey bars, and a key regulatory region is indicated with an asterisk. Data is N=10 for older males, and data for Figure 1D is presented as mean ± SEM.

We subsequently mapped methylation patterns across the 16 kb region and performed differentially methylated region (DMR) analyses to identify loci that were prone to methylation changes following resistance training. Using a 100 bp size sliding window model, with stringent significance thresholds set (e.g. q value < 0.01, differential change of > 5% and with at least 3 contiguous 100 bp windows identified), we identified four DMRs (Figure 1D, Suppl. File 1B). Interestingly, this analysis identified a DMR spanning a larger 500 bp region whose origin mapped to that of the D-loop/control region of the human mtDNA genome (Figure 1D).

The entire control region spans a 1.12 kb locus containing a number of extra regulatory elements including the hyper-variable region (HVR), a tertiary DNA fragment (7S DNA), control elements (Mt5 and Mt3L) and TFAM binding sites (Figure 2a). Crucially, the 1.12 kb locus also contains the light strand promotor (LSP), one of the two heavy strand promotors (HSP1), and the HSP2 promoter resides less than 100 bp away from this region (Figure 2A). Given that we identified a DMR within this control region (Figure 1D), and this region regulates mtDNA replication and transcription, we examined the locus that spans the mtDNA control region as well as the HSP2 region (from 16024 to 650 bp, Figure 2A). In this region, we also identified differentially methylated profiles of the CpGs following resistance training in older participants (Figure 2B). Four of five CpG sites residing within close proximity to either HSP1, HSP2 or LSP showed a significant reduction in methylation (FDR < 0.05, change of > 5%; Figure 2B). This suggests the control region, and in particular the 5’-prime end, is largely hypomethylated following resistance training in older participants.

**Figure 2.**
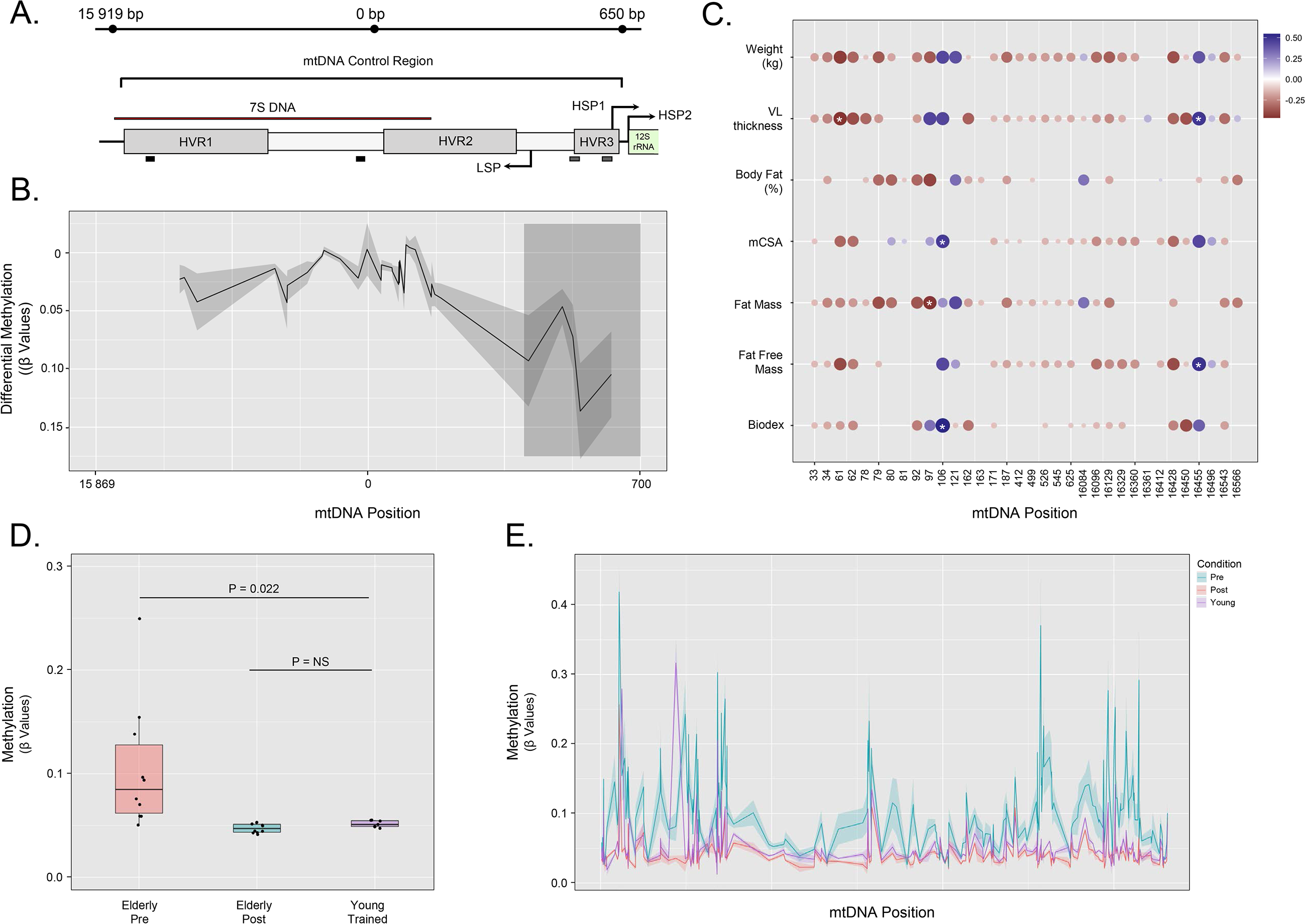
Methylation of the mtDNA regulatory region and correlational analyses with phenotypic variables. Legend: Given that our analyses identified a differentially methylated region (DMR) spanning the known regulatory region of the mtDNA, we further analyzed a 1.12 kb span within this region (panel A). Here, we demonstrated that there was significant differential methylation (FDR < 0.05) in older males with six weeks of training within this region (grey shaded region with 4/5 CpG sites with difference of >5%, panel B). The methylation of CpG sites residing within this locus demonstrated significant correlations with phenotype data (indicated with asterisks) (panel C). Dot coloration represents positive or negative associations, with size and strength of color representing strength of the correlation coefficient. Comparison of mtDNA methylation of younger, trained versus older males prior to training showed a clear hypomethylation (P = 0.022), whereas no such difference is observed with older males following training (panel D). This is further represented across the mtDNA genome highlighting the restoration of the mtDNA methylome to youth like levels via resistance training in older males (Figure 2E). Data is N=10 for all comparisons/correlations of older males, and N=7 for younger trained males. Data for panels B and E are presented as mean ± SEM values.

Associating CpG site methylation data in older participants prior to and following training against phenotype variables, we also identified several significant correlations (Figure 2C; Suppl. File 2). CpG sites residing at positions 61 and 97 of the mtDNA genome showed significant inverse correlations between CpG methylation and vastus lateralis thickness as well as whole-body fat mass, respectively (Figure 2C). These same positions also showed association trends between other phenotype variables within our data sets (albeit not significant; Suppl. File 2). Interestingly, two sites (positions 106 and 16455) in our analyses displayed positive correlations between methylation and phenotype variables. Methylation of site 106 strongly correlated with knee extensor peak torque (P = 0.01) and, to a lesser extent, muscle cross sectional area (P = 0.03). Finally, methylation of position 16455 significantly correlated with both whole-body fat free mass (P = 0.02) and vastus lateralis thickness (P = 0.03).

Next, we compared the methylation profiles of our older males prior to and following training to younger males to ascertain whether training in the older participants restores mtDNA methylation levels to youth like levels. A significant difference was observed between younger and older males prior to training (P = 0.014). Strikingly, however, methylation patterns of younger participants showed no significant difference compared to older participants following training (P > 0.05). Additionally, across the mtDNA genome, a highly comparable methylation profile existed between younger and older males following training (Figure 2E). Collectively, these data suggest resistance training restored the mtDNA methylome in older participants to mimic a more youthful signature.

### Mitochondrial gene expression and protein complexes

The majority of mtDNA genes are transcribed from the HSP2 region. The primary role of HSP1 is to transcribe mitochondrial rRNA genes on the heavy strand. LSP1 transcribes ND6, which is the only coding transcribing gene on the light strand. We therefore undertook qPCR to assess mitochondrial gene expression of MT-CYB (cytochrome B), MT-ND5 (NADH dehydrogenase 5) and CO1 (cytochrome c oxidase subunit I) as well as MT-RNR2 (mitochondrially-encoded 16S rRNA) and light strand ND6 (NADH dehydrogenase 6) mRNA levels. Given TFAM binding sites were located close to the hypomethylated region identified in the DMR analysis above we also measured TFAM gene expression. Resistance training in older participants increased all assayed mitochondrial mRNA targets (MT-CYB, MT-ND6, MT-ND5, and MT-CO1) as well as MT-RNR2 RNA levels (P < 0.05) (Figure 3A), but not TFAM expression (Figure 3B; p > 0.05). We also confirmed no change in TFAM protein levels (P > 0.05; Figure 3C). All of the genes analyzed demonstrated higher levels in the younger compared to older males at both the pre- and post-training time points (P < 0.05 for all targets). Furthermore, citrate synthase activity assays were performed to assess mitochondrial volume (Figure 3D), and no change occurred in older individuals with resistance training (P = 1.00). Skeletal muscle protein levels of electron transport chain complexes were analyzed, and resistance training in older participants increased protein levels of complexes III and IV (P < 0.05) (Figure 3E/F).

**Figure 3.**
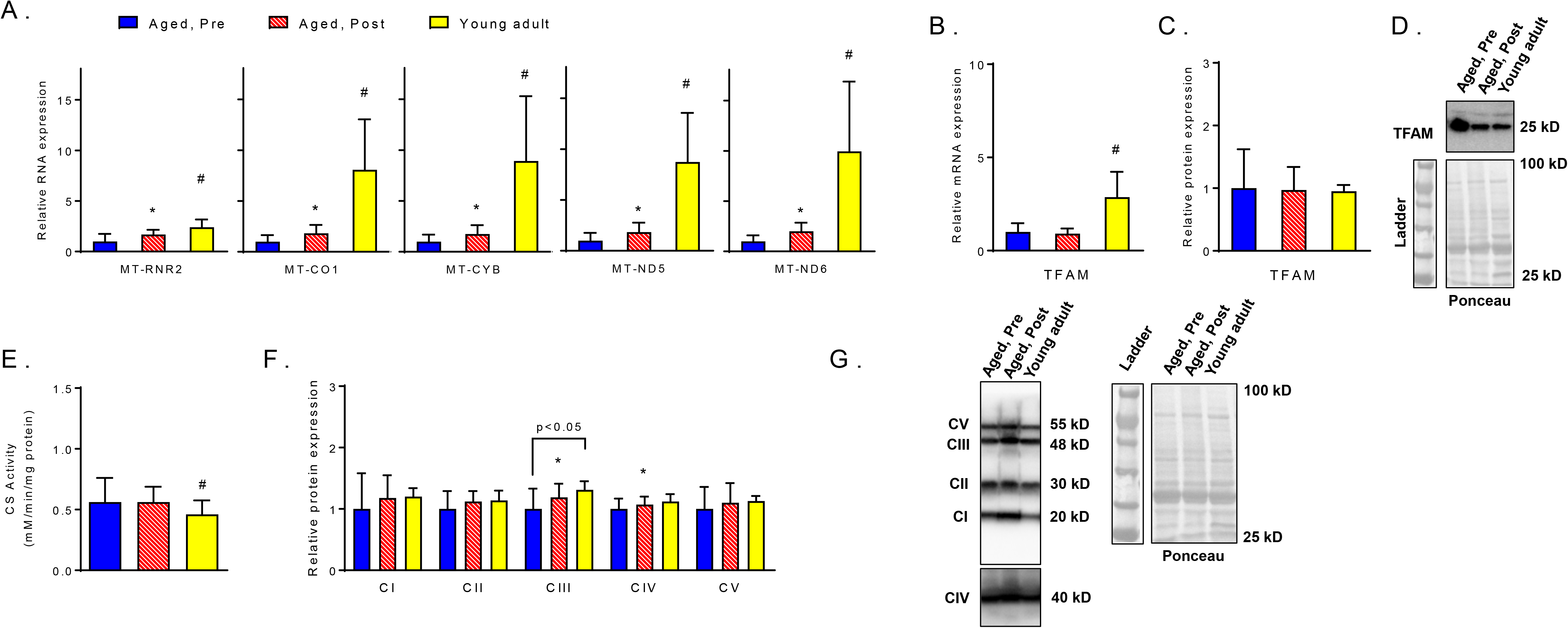
Mitochondrial transcript and marker adaptations with training in older males. Legend: These data represent mitochondrial transcript levels (panel A), TFAM mRNA levels (panel B), TFAM protein levels (panel C), citrate synthase (CS) activity levels (panel E), and protein levels of mitochondrial complexes I-V (panel F). Panels D and G contain representative Western blots for data in panels C and F, respectively. qPCR data contain n=10 older males prior to and following training, and n=7 younger trained males. Citrate synthase activity data contain n=10 older males prior to and following training, and n=6 younger trained males. Western blot contain n=9 older males prior to and following training, and n=6 younger trained males. Symbols: *, indicates increase with training in older males (p<0.05); #, indicates different from younger trained males prior to and following training (p<0.05). Gene abbreviations can be found in-text. All data are presented as means ± SD values.

### Correlation of methylation with mRNA expression and protein levels

We examined associations between CpG methylation patterns and alterations in gene expression to explore the transcriptional consequence of the differentially methylated regions identified. A clear inverse association existed between CpG site methylation (CpG sites residing in our 1.12kb mtDNA loci) and mitochondrial gene expression (Figure 4A). Interestingly, our analysis identified MT-ND6 and MT-RNR2 expression to be the two most commonly inversely associated transcripts (Suppl. Fig 2A and 2B), with CpG site 162 showing the strongest inverse association with MT-RNR2 expression (r = 0.59, P = 0.005) (Figure 4C). Across all analyses, CpG site 16,329, which resides in HVR1, displayed the most consistent negative/inverse correlation between methylation and gene expression (Suppl. File 3). Counterintuitively, but in keeping with correlational analyses performed on our phenotype data sets, CpG site 16455 positively correlated with expression of our analysed gene sets (Figure 4A/B, Suppl. Figure 2A, Suppl. File 3). However, this CpG site is not positioned within any key regulatory locus.

**Figure 4.**
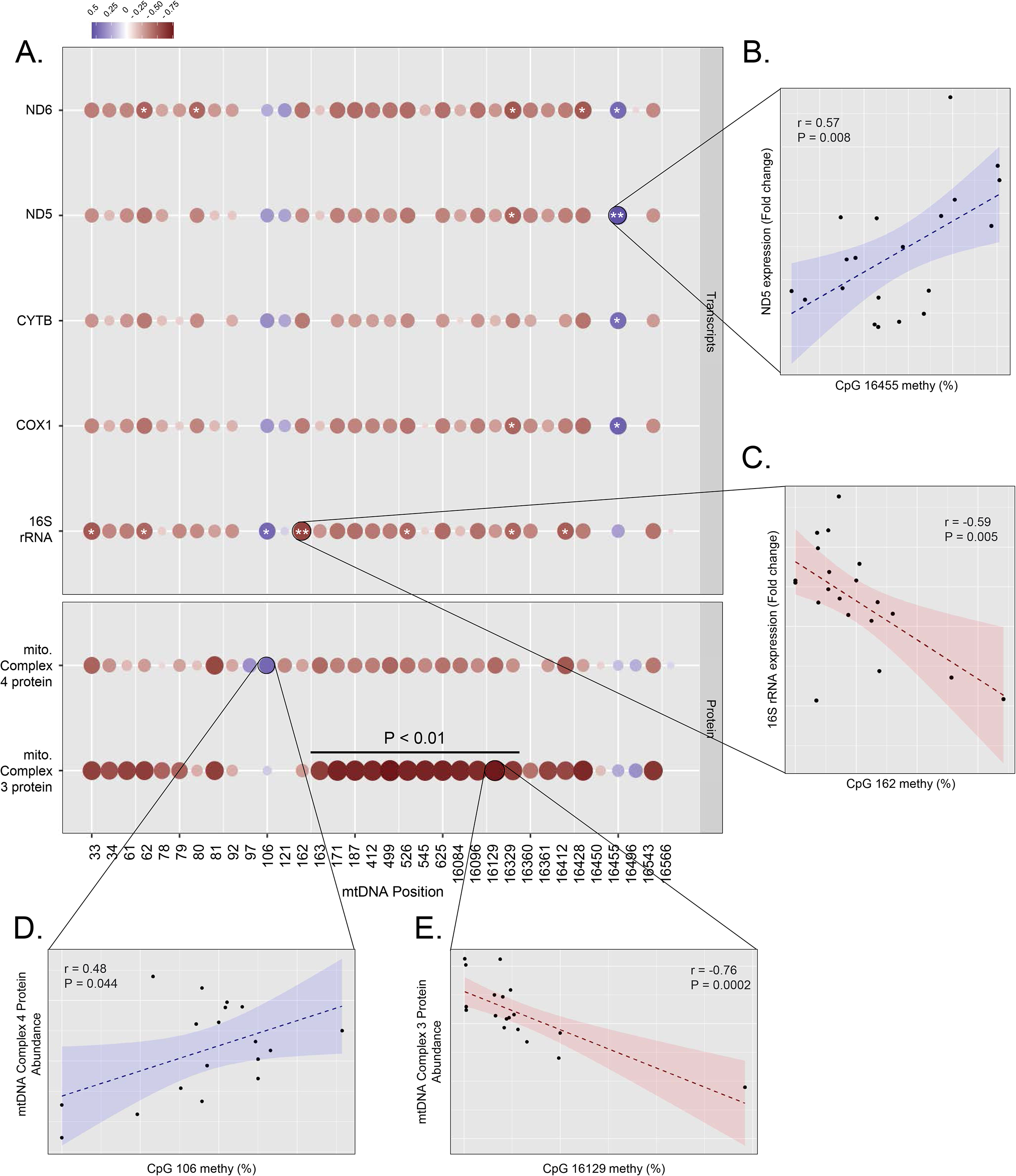
Correlation between CpG methylation, gene expression and protein abundance in older males prior to and following resistance training. Legend: Correlation of methylation in CpG sites residing within the 1.12 kb locus of interest in older males prior to and following training shows significant (* = P < 0.05, ** P<0.01) associations between methylation and gene expression (panel A, facet 1). Coloration of dots represents direction of correlation, with strength of color and size of dot representing the strength of correlation coefficients. Highlighted comparisons demonstrated a positive correlation between CpG 1645 methylation and ND5 gene expression (r = 0.57, P = 0.008), and an inverse correlation between CpG 162 methylation and 16S rRNA expression (r = −0.59, P = 0.005). Correlation of methylation levels in older males prior to and following training with protein abundance of complexes 3 and 4 yielded more striking associations (for all data with relative significance see Suppl. File 4). Of note, CpG sites 163 to 16329 inversely correlated with mtDNA complex 3 protein abundance (P < 0.01), with panel E highlighting the strongest association within (CpG 16129; r = −0.76, P = 0.0002). Positive correlations between these data sets also existed where, for example, CpG 106 methylation positively correlated with complex 4 (panel D; r = 0.48, P = 0.044). Data is N=10 for all comparisons/correlations.

We further correlated methylation with the abundance of complex 3 and 4 proteins (Figure 4A). Strikingly, methylation of CpG sites 163 to 16329 (Figure A) demonstrated a strong inverse association with complex 3 protein abundance (P < 0.001; Figure 4A; Suppl. File 4). CpG site 16129 demonstrated an inverse association (r = −0.76, P = 0.0002; Figure 4E). In keeping with the identification of the positive correlations between CpG methylation of site 106 and phenotypic variables as well as gene expression, we identified a positive correlation (r = 0.48, P = 0.04) between the methylation of this site and complex 4 protein abundance (Figures 4A/D), but such an association was not evident with complex 3 protein abundance.

## DISCUSSION

This study is the first to illustrate that resistance training leads to a robust hypomethylation of the mitochondrial genome in skeletal muscle. Moreover, resistance training seemingly restored mtDNA methylation signatures in older males relative to younger, trained males. Pre- to post training increases in mitochondrial mRNA and rRNA levels in older participants aligned with the observation that the D loop/control region, which regulates mitochondrial transcription and replication, demonstrated increased hypomethylation with training. There were also interesting associations between mtDNA methylation patterns and various phenotypes. These findings are discussed in greater detail below. From a healthy aging perspective, these data continue to suggest resistance training has beneficial effects on certain aspects of mitochondrial physiology.

Skeletal muscle DNA methylation has been reported to increase with aging, and this typically coincides with a decrease in the mRNAs of genes that exist downstream of methylated regions (Blocquiaux et al., 2020; Day et al., 2013; Ling et al., 2007; Ronn et al., 2008; Turner et al., 2020). However, data are lacking in regards to how aging affects the methylation status of the mitochondrial genome in skeletal muscle. D’Aquila et al. (D’Aquila et al., 2015) demonstrated that the methylation of the 12S rRNA region of the mitochondrial genome in PBMCs increases with aging, and 9-year follow-up data illustrate that increased methylation in this region is associated with increased mortality. The current data are in agreement with the findings of D’Aquila and colleagues in that skeletal muscle from older participants prior to training displayed increased mtDNA methylation levels compared to younger, trained participants. Moreover, these methylation patterns coincided with lower mRNA and rRNA levels of various mitochondrial genes as well as lower protein abundances of certain complexes in the older participants. Remarkably, resistance training in older participants decreased mtDNA methylation patterns in certain regions. Although longer-term exercise training has been shown to alter skeletal muscle DNA methylation patterns in younger (Bagley et al., 2020; Seaborne, Strauss, Cocks, Shepherd, O’Brien, Someren, et al., 2018) and middle-aged (Nitert et al., 2012) participants, results from these studies suggest that various genes can be hypo- or hypermethylated. Furthermore, none of these studies interrogated the mtDNA methylation changes given that chip arrays lacking mtDNA probes were utilized. The current findings agree in principle with a recent meta-analysis that examined 16 research studies and concluded that nuclear DNA methylation generally decreases with exercise in older adults (Brown, 2015). Additionally, our data agree in principle with other findings that show resistance exercise evokes nuclear genome hypomethylation in older human skeletal muscle (Blocquiaux et al., 2020). However, our findings strongly extend the current literature given that it is the first to suggest exercise training can lead to the hypomethylation of the mitochondrial genome, and DMR analysis demonstrates this hypomethylation is particularly enriched in important regulatory regions for mtDNA transcription and replication.

Prior to discussing the implications of the mtDNA methylation data, it is important to appreciate how the nuclear and mitochondrial genomes differ. The nuclear genome contains approximately 3 billion base pairs, encodes for just over 20,000 genes, and each gene typically contains one segment of DNA separated by regions of non-coding DNA. mtDNA contains 16,569 base pairs, and encodes for 37 genes including 2 rRNAs, 22 tRNAs and 13 proteins. The mitochondrial genome possesses a heavy strand (H-strand) and light strand (L-strand) where polycistronic RNAs are transcribed from each strand, and subsequently cleaved and processed to yield mitochondrial rRNAs, tRNAs and mRNAs (Shokolenko & Alexeyev, 2017; Taanman, 1999). Two transcription initiation sites exist in a region termed the D-loop. These sites are termed heavy strand promoter 1 (HSP1) and light strand promoter (LSP), and each is where H-strand and L-strand transcription initiation occurs, respectively. Of the mtDNA regions that demonstrated enriched hypomethylation with resistance training in older participants, the most interesting site identified was the D-loop/control region. This finding suggests that resistance training promotes a favorable environment for increased mitochondrial transcription, and aligns with our findings of increased RNA levels H-strand genes (MT-RNR2, MT-CO1, MT-CYB, MT-ND5) and an L-strand gene (MT-ND6) in older individuals following training. While these findings are novel and provocative, it is unclear as to whether the observed methylation and RNA adaptations in older individuals directly facilitated mitochondrial adaptations. In this regard, citrate synthase activity levels, which are strongly associated with mitochondrial volume, remained unaltered with training in older participants. Likewise, only select mitochondrial proteins (specifically, complexes III and IV) were upregulated with training. It is notable, however, that our laboratory has reported 10 weeks of resistance training increases mitochondrial biogenesis in older participants (Lamb, Moore, Mesquita, et al., 2020). Moreover, others have reported that longer-term resistance training (~3 months) increases various aspects of mitochondrial function (e.g., respiration and/or complex activities) in older participants (Holloway et al., 2018; Parise et al., 2005; Robinson et al., 2017). Mitochondrial adaptations involve a coordinated effort between the nuclear and mitochondrial genomes given that most mitochondrial proteins are encoded by the nuclear genome (Parry et al., 2020). Thus, the observed mtDNA methylation changes with resistance training may precede certain longer-term mitochondrial adaptations.

We were also interested in determining whether TFAM mRNA or protein levels were altered in older participants with resistance training given that TFAM is mitochondrial transcription factor that binds to various regions in the D-loop and is critical for stimulating transcription. There were no alterations in TFAM mRNA or protein levels in older participants with resistance training. However, this does not exclude the possibility that TFAM binding to the mitochondrial HSP1 and LSP regions increased during various periods throughout the six-week training protocol. Due to tissue limitations, we were not able to perform assays relevant to assessing this phenomenon. However, certain molecular analyses (e.g., ChIP-qPCR or electrophoretic mobility shift assays) can be performed in the future to address this question.

While we posit that these are novel and exciting findings for the fields of molecular exercise science and muscle aging biology, outstanding questions remain. First, this study is limited to males, and future research is needed to address whether the adaptations observed herein are also observed in females. The inclusion of a younger, trained male cohort was for comparative purposes only. However, it is unknown if resistance training can facilitate the same adaptations in this population as well. Due to tissue limitations, we did not address the mechanism(s) through which mitochondrial demethylation occurred. DNA demethylation (hypomethylation) can occur through the conversion of methylcytosine to hydroxymethylcytosine via ten-eleven translocation (TET) enzymes, and methylation occurs via the de novo methyltransferases (DNMTs) (Wu & Zhang, 2017). Thus, examining whether resistance training either acutely or chronically downregulates mitochondrial DNMT activity and/or upregulates mitochondrial TET enzyme activity is warranted. It is also notable that a recent review by two of the current co-authors provides evidence to suggest that exercise-induced alterations in muscle metabolites can affect enzymes involved with nuclear (and presumably mitochondrial) DNA hypomethylation (Seaborne & Sharples, 2020). Specifically, the authors noted that numerous TCA cycle intermediaries (e.g., FAD/FADH_2_ ratio, alpha-ketoglutarate, succinate, and fumurate levels) can all influence demethylase activity. Given that the TCA cycle occurs in the mitochondrial matrix, a metabolomics approach in isolated skeletal muscle mitochondria prior to and transiently following a resistance exercise bout could provide clues as to whether metabolic perturbations are associated with some of the methylation patterns observed herein. The assayed mitochondrial markers were also limited in scope. In this regard, markers of mitochondrial function (e.g., state III/IV respiration rates with different substrates or complex activities) were not examined, and it is possible that some of these markers also coincided with some of the observed molecular adaptations. Moreover, only enough tissue was available to run biochemical assays on crude muscle lysates rather than isolated mitochondria.

### Conclusions

This is the first study to suggest resistance training in older individuals leads to an appreciable hypomethylation of the mtDNA genome and, specifically, in important regulatory regions. Moreover, observed methylation changes were associated with an increase in various mitochondrial transcripts. Importantly, resistance training restored the mtDNA methylome of older individuals towards profiles observed in younger, trained adults. However, it remains unknown as to whether these events preceded and/or facilitated certain mitochondrial adaptations. Therefore, more research is needed to interpret the significance of these findings.

## EXPERIMENTAL PROCEDURES

### Ethical approval

This study was a secondary analysis of two studies approved by the Institutional Review Board at Auburn University. The first protocol (Protocol # 19-249 MR 1907) involved examining the effects of resistance training with daily peanut protein supplementation or no supplementation on skeletal muscle hypertrophy in untrained, older adults between the ages of 50 to 75 years (NCT04015479). Ten older males from the 6-week cohort (n=5 per group) were examined herein. Two-way repeated measures ANOVAs indicated that none of the body composition or assayed biomarkers were affected by peanut protein supplementation (interaction p-values: lean body mass, p=0.952; vastus lateralis (VL) thickness, p=0.543; knee extensor peak torque, p=0.893; all qPCR and Western blot markers, p>0.200; CS activity, p=0.335). The second protocol involved examining the effects of unilateral resistance training on muscle hypertrophy outcomes in seven previously trained young adult males (Protocol # 19-245 MR 1907). Resting baseline biopsies from these participants were used as a comparator group to the older participants to determine if resistance training rejuvenated the mtDNA methylome.

Inclusion criteria for both studies required participants to abstain from nutritional supplementation (e.g., creatine monohydrate, protein supplements) one month prior to testing. Participants from both studies had to be free of overt cardio-metabolic diseases (e.g., type II diabetes, severe hypertension, heart failure) or conditions that precluded the collection of a skeletal muscle biopsy. All participants provided verbal and written consent to participate in each respective study, and both studies conformed to standards set by the latest revision of the Declaration of Helsinki. Data herein included 10 older male participants (age = 65 ± 7 years old; mean ± SD), and 7 previously trained younger adult males (22 ± 2 years old, self-reported resistance training experience of 5 ± 1 years).

### Resistance training program for older participants

The training program for older males has been previously described (Lamb, Moore, Smith, et al., 2020). Briefly, participants underwent supervised resistance training twice weekly, on non-consecutive days, for six weeks. Each session consisted of five exercises including leg press, leg extensions, lying leg curls, barbell bench press, and cable pull downs. For each exercise, participants performed three sets of 8-12 repetitions to volitional fatigue with at least one minute of rest in between sets. At the end of each set, participants were asked to rate the level of difficulty (0 = easy, 10 = hard). If values were below 7, weight was added to increase effort for the next working set. If values were 10, or the participant could not complete the set, weight was removed prior to the next working set. Participants were encouraged to be as truthful as possible when assessing difficulty. The intent of this training method was to challenge participants where perceived exertion after each set was between a 7-9 rating. This method allowed us to ensure that training effort was maximized within each training session, and that the participants were successfully implementing progressive overload in an individualized fashion.

### Testing sessions

For younger and older participants, the testing sessions described below occurred during morning hours (05:00–09:00) following an overnight fast. For older males, Pre-testing occurred ~2-5 days prior to the first day of resistance training, and Post-testing occurred 72 hours following the last training bout. The younger males performed all of the same tests described above between 05:00-11:00.

Prior to testing batteries, participants submitted a urine sample (~5 mL) to assess urine specific gravity (USG) using a handheld refractometer (ATAGO; Bellevue, WA, USA). USG in all participants were <1.020 indicating sufficient hydration (American College of Sports et al., 2007). Height and body mass were assessed using a digital column scale (Seca 769; Hanover, MD, USA), and values were recorded to the nearest 0.1 kg and 0.5 cm, respectively. Participants then had their bone-free lean/soft tissue mass (LSTM) and fat mass determined by a full-body dual-energy x-ray absorptiometry (DXA) scan (Lunar Prodigy; GE Corporation, Fairfield, CT, USA). The same investigator completed all DXA scans. According to previous data published by our laboratory (Kephart et al., 2016), the same-day test-calibrate-retest reliability on 10 participants produced an intra-class correlation coefficient (ICC) of 0.998 for LSTM. After DXA scans, a cross-sectional image of the right thigh at 50% of the femur length was acquired using a peripheral quantitative computed tomography (pQCT) scanner (Stratec XCT 3000, Stratec Medical, Pforzheim, Germany). Scans were acquired using a single 2.4 mm slice thickness, a voxel size of 0.4 mm and scanning speed of 20 mm/sec. Images were analyzed for total muscle cross-sectional area (mCSA, cm^2^) using the pQCT BoneJ plugin freely available through ImageJ analysis software (NIH, Bethesda, MD, USA). All scans were performed and analyzed by the same investigator, and the ICC was previously determined for mCSA to be 0.990 (*unpublished data*). Following pQCT assessments, right leg vastus lateralis ultrasound assessments were performed using a 3-12 MHz multi-frequency linear phase array transducer (Logiq S7 R2 Expert; General Electric, Fairfield, CT, USA) to determine muscle thickness. Participants were instructed to stand and displace bodyweight more to the left leg to ensure the right leg was relaxed. Measurements were standardized by placing the transducer at the midway point between the inguinal crease and proximal patella. The same technician performed all ultrasounds. According to previous data from our laboratory, the 24-hour test-retest reliability for muscle thickness assessment on 11 participants resulted in an ICC of 0.983.

Right leg vastus lateralis muscle biopsies were then obtained with a 5-gauge needle as previously described (Kephart et al., 2015). Following biopsies, tissue was rapidly teased of blood and connective tissue, wrapped in pre-labeled foils, flash frozen in liquid nitrogen, and subsequently stored at −80°C for further molecular analyses.

In older males, right leg knee extensor peak torque testing occurred ~1-3 days prior to the muscle biopsy at the pre time point, whereas this test occurred approximately 30 minutes prior to the biopsy at the post-test time point. In younger males, this test occurred approximately 10 minutes following the biopsy. During testing, participants were fastened to an isokinetic dynamometer (Biodex System 4; Biodex Medical Systems, Inc., Shirley, NY, USA). Each participant’s knee was aligned with the axis of the dynamometer, and seat height was adjusted to ensure the hip angle was approximately 90°. Prior to peak torque assessment, each participant performed a warmup consisting of submaximal to maximal isokinetic knee extensions. Participants then completed five maximal voluntary isokinetic knee extension actions at 60°/s. Participants were provided verbal encouragement during each contraction. The isokinetic extension resulting in the greatest value for peak torque was used for analyses.

### Molecular analyses of skeletal muscle

#### DNA isolation

Muscle samples stored in foils were removed from −80°C and placed on a liquid nitrogen-cooled ceramic mortar. Tissue was crushed using a ceramic pestle, and ~10 mg was obtained for DNA isolation using the commercially available DNeasy Blood & Tissue Kit (Qiagen; Venlo, The Netherlands; catalog #: 69504) as per the manufacturer’s recommendations. DNA pellets were reconstituted in a buffer provided by the kit, and concentrations were determined in duplicate at an absorbance of 260/280 nm (1.81 ± 0.08) using a desktop spectrophotometer (NanoDrop Lite; Thermo Fisher Scientific; Waltham, MA, USA). DNA was then shipped to a commercial vendor (EpiGentek Group Inc.; Farmingdale, NY, USA) for RRBS as described below.

#### DNA bisulfite conversion and RRBS

Samples were received by the commercial vendor on dry ice, and were subjected to enzymatic digestion (MSP1 + TaqI); specifically, 300 ng of DNA from each participant was digested for 2 hours with MSP1 enzyme (20U/sample) at 37°C followed by 2 hours with TaqαI (20U/sample) at 65°C. Digested DNA <300 base pair fragments were collected for bisulfite treatment, and bisulfite conversion was performed with the Methylamp DNA Bisulfite Conversion Kit (Epigentek; catalog #: P-1001). The efficiency of bisulfite-treated DNA was determined by real-time PCR using two pairs of primers where the first pair targeted bisulfite-converted beta-actin (BACT), and the second pair targeted unconverted Glyceraldehyde-3-Phosphate Dehydrogenase (GAPDH) for the same bisulfite-treated DNA samples. Library preparation then ensued, and Bioanalyzer QC and KAPA library quantification were performed thereafter. Sample libraries (20 nM) were subjected to multiplex next generation sequencing using an Illumina HiSeq4000 (Illumina Inc.; San Diego, CA, USA). Quality control on raw reads was performed using FASTQC, version 0.11.8 (http://www.bioinformatics.bbsrc.ac.uk/projects/fastqc/), and an HTML report was generated for each data set. Quality and adapter trimming was performed on the raw reads using Trim Galore, version 0.5.0 (http://www.bioinformatics.babraham.ac.uk/projects/trim_galore/). Trim Galore performs the following trimming steps: i) low-quality read removal (Sanger Phred score of 20 or lower), ii) trimming of the 3’ Illumina adapter (any signs of AGATCGGAAGAGC), and iii) removal of trimmed reads shorter than 20 bp. Trimmed reads were mapped to the UCSC homo sapiens (human) genome sequence (version GRCh38) using a methylation-aware mapper, bismark, version 0.203.0 (Krueger & Andrews, 2011). Bismark utilizes Bowtie, version 2.2.5 (Langmead, Trapnell, Pop, & Salzberg, 2009), with the option “--directional” for targeted bisulfite sequencing libraries and option “--pbat” for post-bisulfite prepared RRBS libraries. For each sample, a summary HTML report was generated, which included alignment and cytosine methylation statistics. Samtools, version 0.1.9 (Li et al., 2009), was utilized to sort the SAM file produced by bismark and remove the duplicate reads due to PCR amplification. Methylation information was extracted from the final bismark mapping result at the base resolution where a minimal read coverage score of 10 and minimal quality score of 20 at each base position are applied. For all comparisons utilized in older adult trained versus untrained data sets, this meant 254 CpG sites were covered within the mtDNA. For inclusion of the younger trained adult data set, this number reduced to 253 CpG sites in the mtDNA. The resulting CpG sites were filtered based on coverage and merged for comparative analysis using MethylKit package (https://github.com/al2na/methylKit) in R (version 4.0.3). Only CpG sites that were covered in all participants were merged. Principle Component Analysis plots were subsequently performed to determine group-level quality control. Differentially Methylated Regions (DMRs) and Differentially Methylated CpGs (DMCs) were processed in a similar manner. However, for DMR analysis, data sets were first chunked into 100bp windows with a step size of 100bp. Differential analysis was then performed using MethylKits *calculateDiffmeth* function and logistic regression to calculate differential P values which were then transformed to Q values using the SLIM method (Wang, Tuominen, & Tsai, 2011), and DMRs/DMCs were extracted.

#### RNA isolation with Trizol and targeted qPCR

Approximately 10 mg of muscle was placed in 500 μl of Ribozol (Ameresco; Solon, OH, USA), and RNA isolation proceeded following the manufacturer’s instructions. RNA concentrations were determined in duplicate using a NanoDrop Lite (Thermo Fisher Scientific), and cDNA (2 μg) was synthesized using a commercial qScript cDNA SuperMix (Quanta Biosciences; Gaithersburg, MD, USA). Real-time qPCR was performed in a thermal cycler (Bio-Rad Laboratories; Hercules, CA, USA) using SYBR green-based methods and gene-specific primers were designed with publically-available software (Primer3Plus; Cambridge, MA, USA). For all primer sets, pilot qPCR reactions and melt curves indicated that only one amplicon was present. The forward and reverse primer sequences of all genes are listed in Table 2. Fold change values were performed using the 2^∆∆-Cq^ method where 2^∆-Cq^ = 2^(housekeeping gene (HKG) Cq - gene of interest Cq), and 2^∆∆-Cq^ (or fold change) = (2^∆Cq^ value/2^∆Cq^ average of Pre values). GAPDH was used as the reference/HKG gene, and GAPDH Cq values were stable with training in the older participants (pre: 27.85 ± 1.28, post: 27.84 ± 1.25).

**Table 2.**
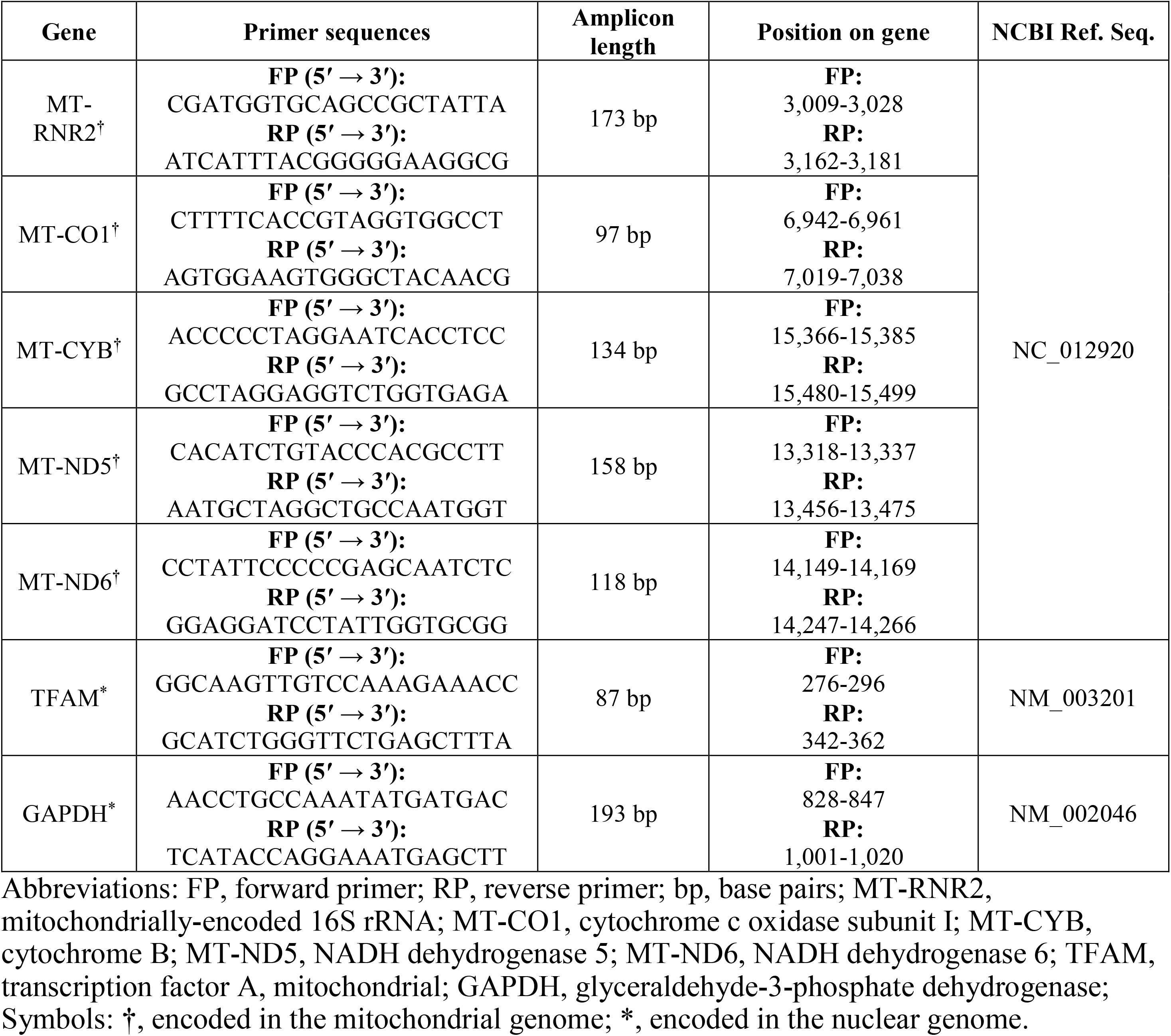
qPCR primers

#### Western blotting (mitochondrial complexes and TFAM protein expression)

Muscle stored in foils were removed from −80°C and placed on a liquid nitrogen-cooled ceramic mortar. Tissue was crushed using a ceramic pestle, and ~20 mg was placed in 1.7 mL microcentrifuge tubes prefilled with general cell lysis buffer (25 mM Tris, pH 7.2, 0.5% Triton X-100, 1x protease inhibitors). Samples were homogenized on ice using hard-plastic pestles, and centrifuged at 1,500 g for 10 minutes at 4°C. Supernatants were collected and placed in new 1.7 mL microtubes on ice. Supernatant protein concentrations were determined using a commercially available BCA kit (Thermo Fisher Scientific; Waltham, MA, USA) according to manufacturer’s instructions. Afterwards, supernatants were prepared for Western blotting using 4x Laemmli buffer and distilled water (diH_2_O) at a concentration of 1 μg/μL, and denatured for 5 minutes at 100°C prior to being frozen at −80°C until Western blotting. On the day of Western blotting, prepared samples (15 μL) were pipetted onto gradient SDS-polyacrylamide gels (4%–15% Criterion TGX Stain-free gels; Bio-Rad Laboratories), and electrophoresis commenced at 180 V for 50 minutes. Following electrophoresis, proteins were transferred to pre-activated PVDF membranes (Bio-Rad Laboratories) for two hours at 200 mA. Gels were then Ponceau stained for five minutes, washed with diH_2_O for one minute, dried for one hour, and digitally imaged with a gel documentation system (ChemiDoc Touch; Bio-Rad Laboratories). Following Ponceau imaging, membranes were re-activated in methanol, blocked with nonfat milk for one hour (5% w/v diluted in Tri-buffered saline with 0.1% Tween 20, or TBST), washed three times in TBST only (5 minutes per wash). Membranes were then incubated for 24 hours with the following antibodies (1:1000 v/v dilution in TBST): i) mouse anti-human OxPhos cocktail (Abcam; Cambridge, MA, USA; catalog#: ab110411), ii) rabbit anti-human TFAM (Abnova; Taipei, Taiwan; catalog #: H00007019-D01P), and iii) COX IV (Cell Signaling Technology; Danvers, MA, USA; Cat# 4850). Following primary antibody incubations, membranes were washed three times in TBST only (5 minutes per wash), and incubated for one hour with horseradish peroxidase-conjugated anti-mouse or anti-rabbit IgG (Cell Signaling Technology; catalog #’s: 7076 and 7074). Membranes were then washed three times in TBST only (five minutes per wash), developed using chemiluminescent substrate (EMD Millipore; Burlington, MA, USA), and digitally imaged using a gel documentation system (ChemiDoc Touch; Bio-Rad Laboratories). Raw target band densities were obtained using associated software (Image Lab v6.0.1; Bio-Rad Laboratories), and these values were divided by Ponceau densities at 25-100 kD. Target/Ponceau density ratios were then divided by the grand mean of older participants at the Pre time point in order to obtain relative protein expression values. For Western blotting, 9 of 10 older participants and 6 of 7 younger males were assayed due to tissue limitations.

#### Determination of muscle citrate synthase activity

Muscle citrate synthase activity levels were determined in duplicate on supernatants obtained from muscle described in the Western blotting section; notably, these methods are similar to previous methods used by our laboratory (Haun et al., 2019; Roberts et al., 2018). The assay principle is based on the reduction of 5,50-dithiobis(2-nitrobenzoic acid) (DTNB) at 412 nm (extinction coefficient 13.6 mmol/L/cm) coupled to the reduction of acetyl-CoA by the citrate synthase reaction in the presence of oxaloacetate. Briefly, 12.5 μg of skeletal muscle protein obtained from supernatants was added to a mixture composed of 0.125 mol/L Tris–HCl (pH 8.0), 0.03 mmol/L acetyl-CoA, and 0.1 mmol/L DTNB. All duplicate reactions occurred in 96-well plates, reactions were initiated by the addition of 5 μL of 50 mmol/L oxaloacetate per well, and the absorbance change was recorded for 60 seconds in a spectrophotometer (Synergy H1; BioTek; Winooski, VT, USA). Again, 6 of 7 younger males were assayed due to tissue limitations.

### Statistics

Phenotype and select molecular data were compared from pre to post training in older males using dependent samples t-tests. Additionally, comparisons of data from older males at these time points were made to younger males using independent samples t-tests. Methylation data were analyzed from pre- to post-training in older males and between older and younger participants using a variety of statistical methods that are described in greater detail in the results section. Select dependent variables were also correlated using Pearson’s correlation coefficients. All data herein are presented in figures and tables as means ± standard deviation values unless stated otherwise, and statistical significance was set at p < 0.05.

## ACKNOWLEDGEMENTS

Participant compensation costs were funded by a grant provided by the Peanut Institute Foundation (Albany, GA, USA) to A.D.F., M.D.R., and K.C.Y. A.P.S was supported by the Norwegian School of Sport Sciences and by The Research Council of Norway (grant number 314157). Assay and APC costs were provided through discretionary laboratory funds from M.D.R.

## AUTHOR CONTRIBUTIONS

B.A.R., A.P.S., R.A.S., and M.D.R. primarily drafted the manuscript and constructed figures. B.A.R., J.S.G., P.H.C.M., S.C.O., C.G.V., D.A.L., K.C.Y., A.N.K., and M.D.R. were involved in critical aspects of the study in regards to data collection and analyses. D.G.C., S.C.F., A.D.F. provided critical assistance in manuscript preparation. All authors edited the manuscript, and all authors approved the final submitted version.

## DATA AVAILABILITY STATEMENT

Several raw data files have been uploaded as supplementary files. Other files can be obtained by emailing the corresponding author.

